# Tandem histone-binding domains enhance the activity of a synthetic chromatin effector

**DOI:** 10.1101/145730

**Authors:** Stefan J. Tekel, Daniel Vargas, Lusheng Song, Joshua LaBaer, Karmella A. Haynes

## Abstract

Fusion proteins that specifically interact with biochemical marks on chromosomes represent a new class of synthetic transcriptional regulators that decode cell state information rather than DNA sequences. In multicellular organisms, information relevant to cell state, tissue identity, and oncogenesis is often encoded as biochemical modifications of histones, which are bound to DNA in eukaryotic nuclei and regulate gene expression states. We have previously reported the development and validation of the “Polycomb-based transcription factor” (PcTF), a fusion protein that recognizes histone modifications through a protein-protein interaction between its polycomb chromodomain (PCD) motif and trimethylated lysine 27 of histone H3 (H3K27me3) at genomic sites. We demonstrated that PcTF activates genes at methyl-histone-enriched loci in cancer-derived cell lines. However, PcTF induces modest activation of a methyl-histone associated reporter compared to a DNA-binding activator. Therefore, we modified PcTF to enhance its target affinity. Here, we demonstrate the activity of a modified regulator called Pc_2_TF, which has two tandem copies of the H3K27me3-binding PCD at the N-terminus. Pc_2_TF shows higher affinity for H3K27me3 *in vitro* and shows enhanced gene activation in HEK293 cells compared to PcTF. These results provide compelling evidence that the intrinsic histone-binding activity of the PCD motif can be used to tune the activity of synthetic histone-binding transcriptional regulators.

## INTRODUCTION

The discovery of histone post-translational modifications (PTMs) and the peptides that specifically interact with these marks has enabled scientists and cell engineers to manipulate chromatin, the DNA-protein structure that regulates gene expression states in eukaryotic cells. Structure-based models have informed targeted knock-down of chromatin subunits and the rational design of low molecular weight inhibitor compounds (reviewed in [1]). DNA-binding domains fused with structural chromatin proteins and histone-modifying enzymes have been used to generate ectopic chromatin conformations at specific loci [2,3]. Until recently, scientists have not yet leveraged PTM-binding peptides from natural effector proteins to “read” the rich biological information encoded in histone marks in living cells. Peptides that recognize specific histone PTM signals are an essential requirement for synthetic systems that integrate epigenetic regulatory signals. In order to use PTM-binding peptides in synthetic fusion proteins, the peptides must be portable, that is, maintain their intrinsic function within a new protein sequence. Early studies established important foundational knowledge by demonstrating that the interaction of the chromodomain motif (CD) with trimethylated histone H3 lysine 27 (H3K27me3) is an intrinsic activity that is maintained by the CD in the context of recombinant, fusion proteins [4,5]. Other protein folds including the bromodomain (BRD) and plant homeodomain finger (PHD) function as isolated peptides [6–8] and within fusion proteins [7,9] to specifically interact with acetylated histone lysines (BRD) and H3K4me3 (PHD).

We constructed the Polycomb-based Transcription factor (PcTF) using a histone PTM-binding motif from the natural protein CBX8 [10]. The CBX8 effector protein binds to histone H3 trimethylated at lysine 27 (H3K27me3) through its N-terminal Polycomb chromodomain (PCD) and establishes a silenced transcriptional state. Expression of PcTF, an artificial transcriptional activator with an N-terminal PCD, mCherry tag, and C-terminal VP64 activation domain, led to increased expression of H3K27me3-enriched genes in three different cancer-derived cell lines [10,11]. These results show promise for designing transcription factors that can read chromatin marks to rewire aberrant epigenetic programming. However, binding affinities observed *in vitro* for isolated PCD domains is poor, reported as 5 - 165 μM [12,13], compared to DNA-binding domains with target affinities in the pico-to nanomolar range such as TALEs (~3 - 220 nM), [14], Zinc Fingers (~0.01 - 16 nM) [15,16], and CRISPR/Cas (~0.5 nM) [17]. In other work, we observed stronger gene upregulation when mCherry-VP64 was targeted to a promoter via a Gal4 DNA binding domain compared to the PCD histone binding domain [18]. PcTF-mediated gene activation is dose-dependent [11], and high PcTF expression levels are required for optimal activity. This limits the usefulness of PcTF for therapeutic applications where barriers to delivery severely limits the number of proteins that ultimately reach the nuclei of target cells. Although pharmacokinetic barriers to DNA and protein delivery *in vivo* are not trivial, increasing the intracellular performance of PcTF is a critical step for advancing this technology towards clinical use.

Here, we extend our investigation of the utility of the PCD motif as a protein design tool by testing its ability to confer multivalency to a synthetic regulator. In this report, multivalency is defined as the engagement of a single protein (monomer) with more than one histone PTM (reviewed in [1,3,19]). Multivalent chromatin proteins can engage adjacent PTMs within a single histone tail, such as K4me3 and R8me2 on histone H3 bound by Spindlin1 [20], or K5ac and K12ac on histone H4 bound by TAF_ǁ_250 [21]. PTM targets can also reside on two distinct histone tails, such as H4K16ac and H3K4me3 bound by BPTF [7]. Dual recognition of histone PTMs is accomplished by tandem protein motifs within the histone-binding protein. Studies have shown that tandem motifs thermodynamically enhance binding affinity and specificity [22]. In order to compensate for the modest affinity of the CBX8 PCD [12] for its target, we added a second copy of H3K27me3-binding PCD to the N-terminus of PcTF to create Pc_2_TF. Here, demonstrate that Pc_2_TF shows higher affinity for H3K27me3 in *vitro.* This activity corresponds with enhanced activation of a H3K27me3-repressed gene in cultured cells.

## RESULTS

### Design of a bivalent synthetic chromatin-based transcriptional regulator

We designed the Pc_2_TF protein to simultaneously recognize two copies of the histone posttranslational modification H3K27me3. The Polycomb chromodomain motif (PCD) consists of three β strands packed against a C-terminal α helix, and a hydrophobic pocket formed by three aromatic residues that interact with a methyl-lysine sidechain [13,23] (Fig. 1A). The arrangement of histones within the nucleosome octamer suggests that Pc_2_TF might bind adjacent trimethylated H3K27 residues. A single nucleosome includes eight individual histone proteins. The central tetramer contains two copies of histone H3 and H4. The H3 proteins are oriented in *cis* so that the unfolded N-terminal tails protrude away from the nucleosome in the same direction [24] (Fig. 1B). One or both H3 tails [25] can become trimethylated at lysine 27 by the enzyme enhancer of zeste (EZH) [26]. Therefore, tandem PCDs in the multivalent protein Pc_2_TF might interact with two histone post translational modifications (PTMs) in a single nucleosome (Fig. 1B) or single PTMs on adjacent nucleosomes.

**Figure 1:**
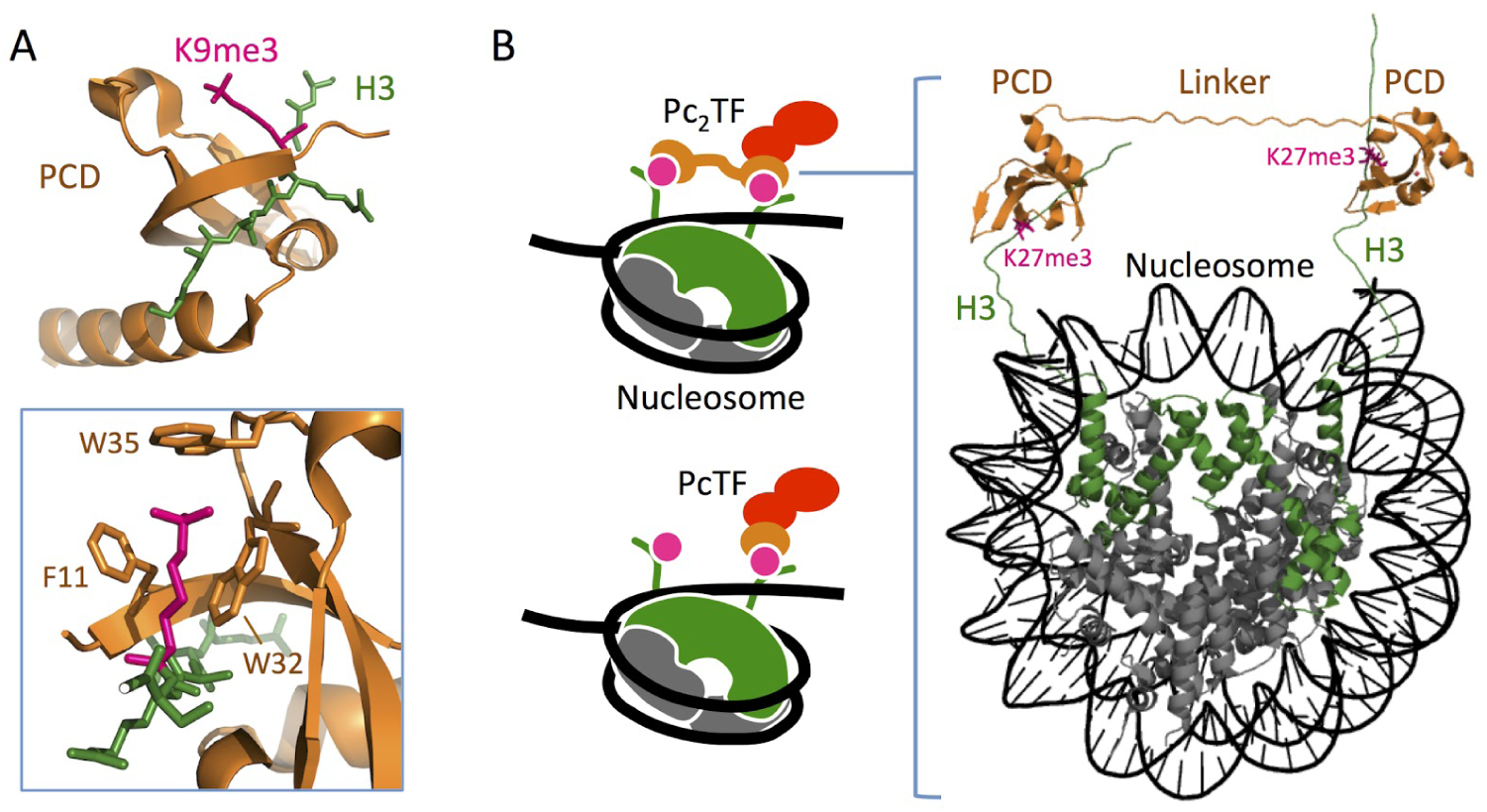
3-D model layout to show the plausibility of Pc_2_TF binding to adjacent H3K27me3 marks. (A) PCD (CBX8) in complex with trimethyl lysine (PDB 3I91) [27]. Three residues form a hydrophobic cage and surround the Kme3 moiety (inset). (B) H3K27me3 recognition by synthetic fusion proteins that carry a single or tandem PCD domains (PcTF and Pc_2_TF, respectively). The 3D rendering was composed in the PyMOL Molecular Graphics System, Version 1.3 Schrödinger, LLC (https://www.pymol.org/) using data for CBX8/H3K9me3 (PDB 4X3K) [28], and a whole nucleosome assembly (PDB 5AV8) [28,29] from the Protein Data Bank.

To identify a linker that allowed optimal binding with H3K27me3, we used an *in vitro* expression and ELISA procedure to test four Pc_2_TF variants. Different lengths and physical characteristics were explored by using flexible glycine-serine linkers [30] and rigid alpha-helical [31,32] linkers. Glycine and serine, amino acids with small side chains, have been used in a wide range protein engineering applications to build flexible linker peptides that have minimal interference with the function of tethered proteins [33]. It has been proposed that flexible, floppy linkers do not support the maximum distance between the tethered peptides [34]. Therefore, rigid linkers might perform better by stabilizing the distance between PCDs to support interactions with neighboring K27me3 moieties. The Pc_2_TF constructs included two tandem copies of the PCD separated by one of four linkers: flexible (GGGGS)_4_, long flexible (GGGGS)_16_, rigid (EAAAR)_4_, and long rigid (EAAAR)_16_. Based on a simplified layout of the interacting components (PCDs and a nucleosome carrying two H3K27me3 modifications) (Fig. 1B), we predicted that 20 amino acids would provide sufficient length for adjacent PCDs to bind simultaneously. The 80 amino acid linkers were used to determine the impact of increased spacing between PCDs.

Recombinant fusion proteins were produced using an efficient, cost-effective bacterial cell-free transcription translation (TXTL) system [35]. We used cell-free expression at the initial prototyping stage to eliminate time-consuming steps involved in *E. coli* over-expression and protein purification. Open reading frames for Pc_2_TF variants and a control protein with no binding domain (Pc_Δ_TF) were cloned into a pET28 vector (Fig. 2A) and added into TXTL solution with a σ70-T7 RNA Polymerase-expressing plasmid. Real-time detection of mCherry fluorescence in a Roche thermal cycler confirmed expression of recombinant proteins. For ELISA, TXTL products were added to microwells in which different C-terminal biotinylated histone peptides (H3 a.a. 21-44) were tethered via pre-coated neutravidin. Significant relative binding over background (unmodified K27 and K27ac) was detected for variants that contained the flexible (GGGGS)_4_, long flexible (GGGGS)_16_, and long rigid (EAAAR)_16_ linkers (Fig. 2C). The implications of these results are discussed in the Conclusions. The flexible (GGGGS)_4_ linker conferred the highest level of binding in this assay. Therefore, we used this variant in subsequent experiments to determine the impact of bivalency on the activity of synthetic, histone-binding effectors.

**Figure 2.**
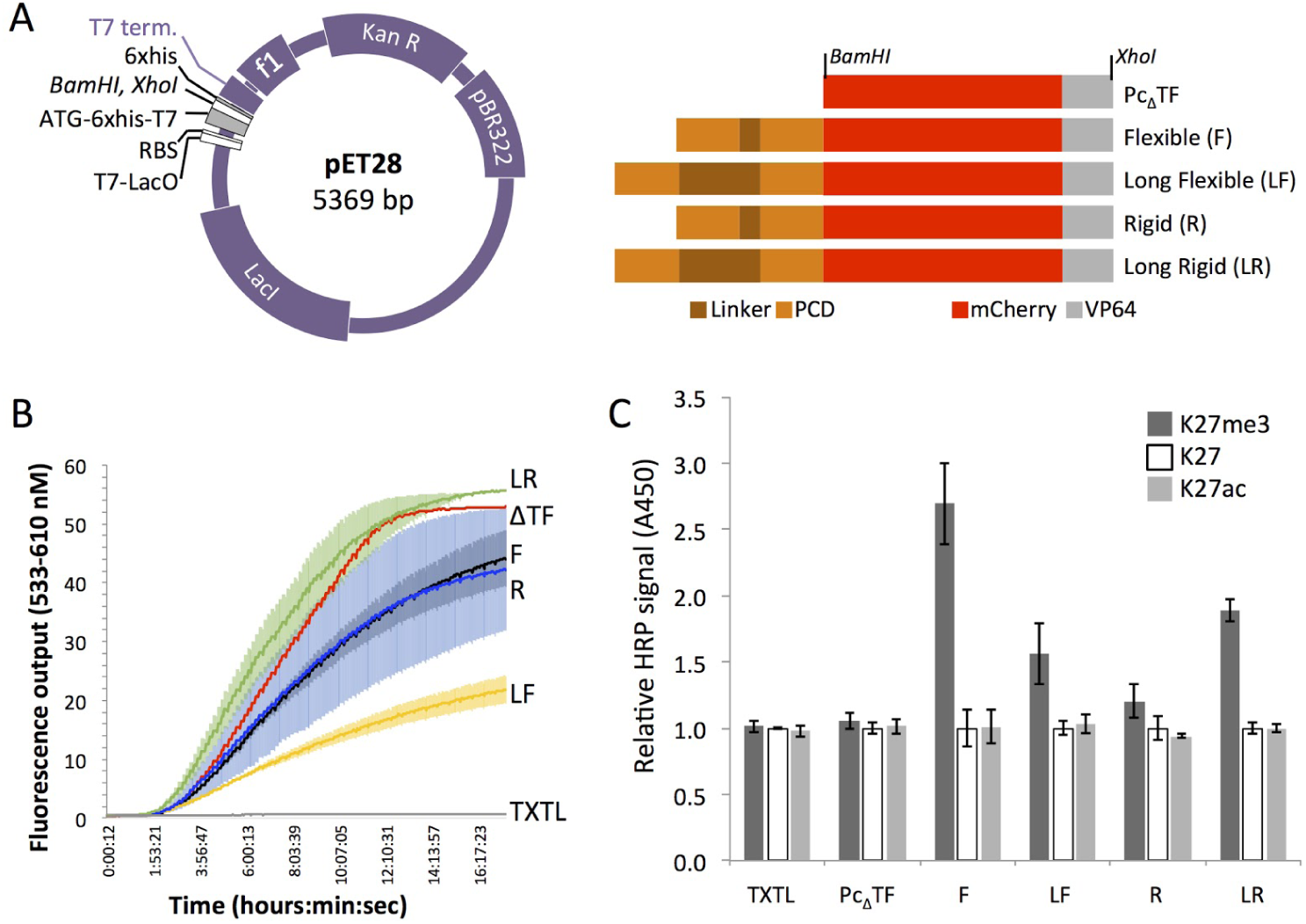
ELISA was used to determine binding affinity and specificity of Pc_2_TF variants that were expressed in a bacterial cell-free expression system. (A) Map of the expression vector and open reading frames (ORFs). Fusion-encoding ORFs were cloned in the pET28 vector at *BamHI* and *XhoI.* (B) Real-time detection of mCherry fluorescence was used to determine expression of recombinant protein in TXTL in a 96-well PCR plate in a Roche thermal cycler. n = 1 for Pc_Δ_TF and TXTL without DNA, n = 3 for others, shaded regions = SDM. (C) The bar chart shows binding of TXTL-expressed fusion proteins (or plasmid-free blank TXTL) with trimethyl-K27, unmodified, or off-target (acetyl-K27) histone H3 peptides. Values for each TXTL sample are normalized to unmodified H3 (n = 3, bars = SDM).

### Bivalent Pc_2_TF shows enhanced H3K27me3 binding compared to PcTF *in vitro*

In ELISA experiments, the bivalent PCD fusion showed greater binding with H3K27me3 peptides compared to the single-PCD fusion protein. The LacI-encoding pET28 vector (Fig. 2A) enabled IPTG-regulatable expression in *E. coli,* which was used to initiate high-yield production once the population reached log growth phase. Native polyacrylamide electrophoresis (PAGE) of lysates from IPTG-treated and untreated *E. coli* showed inducible production of the proteins at roughly the expected sizes: 37, 44, and 52 kilo Daltons for Pc_Δ_TF, PcTF, and Pc_2_TF respectively. Nickel-NTA column-purified proteins were soluble in 1x phosphate buffered saline (PBS). The visible red hue under white light, which is typical of the mCherry protein [36], indicated proper protein folding.

50 nM of purified PcTF, Pc_2_TF, or Pc_Δ_TF was tested for interaction with different C-terminal biotinylated histone peptides as described for the TXTL-expressed proteins. PcTF and Pc_2_TF showed significant binding with H3K27me3 peptides compared to the control protein Pc_Δ_TF (Fig. 3C). Lack of binding with unmodified H3 or H3K27ac indicates that the interaction with H3K27me3 is specific. Pc_2_TF binding showed a 3-fold increase over PcTF. Therefore, at 50 nM paired PCD motifs within the Pc_2_TF protein confer significantly enhanced binding.

**Figure 3.**
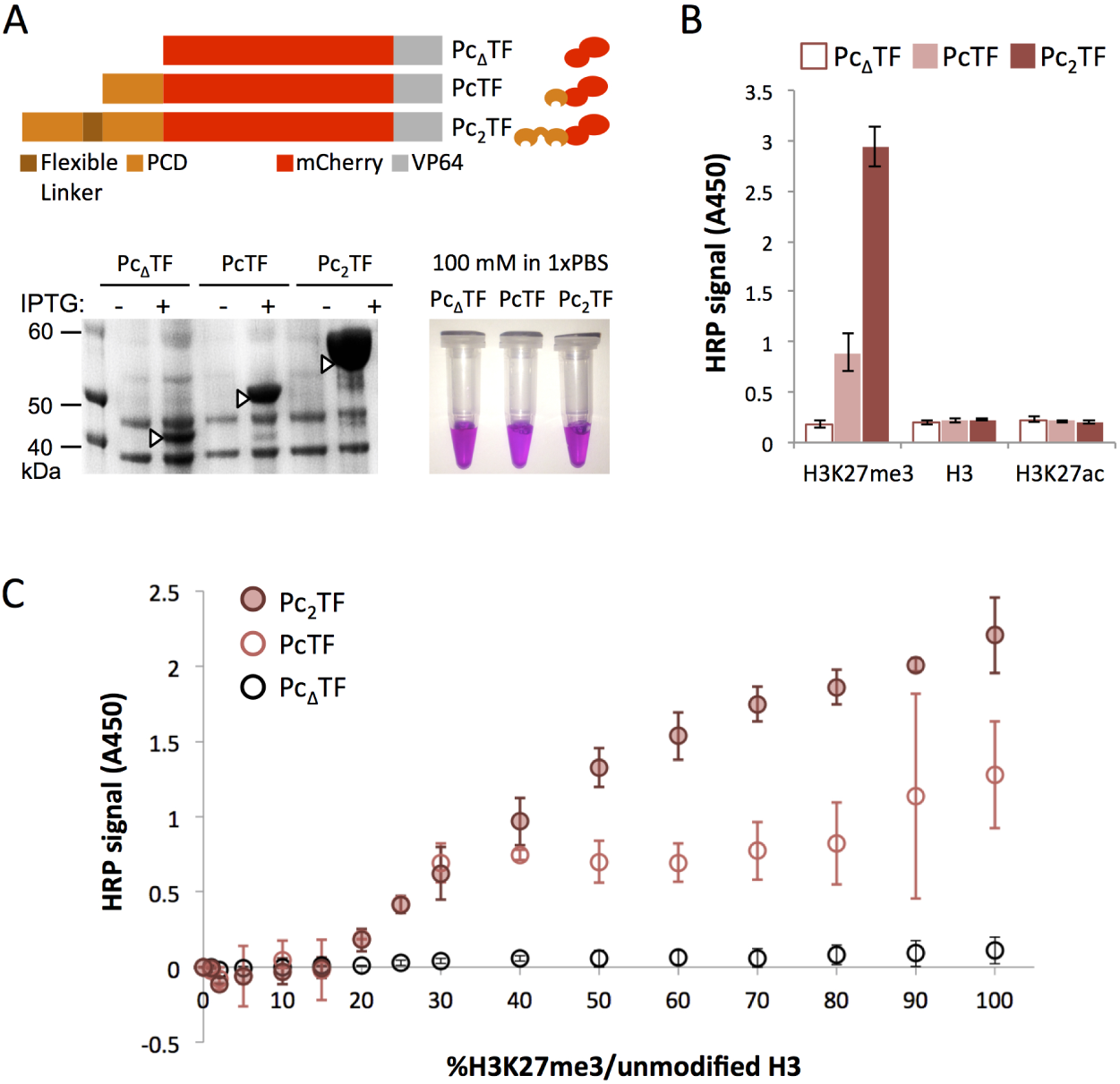
A bivalent PCD fusion peptide shows enhanced H3K27me3 binding *in vitro.* (A) For high-yield expression, *E. coli* was transformed with pET28 plasmids encoding Pc_Δ_TF (negative control), PcTF (single PCD), and the Pc_2_TF containing the flexible linker (GGGGS)_4_. Native polyacrylamide gel electrophoresis (PAGE) of over-expressed proteins purified from *E. coli.*(C) ELISA was used to detect interaction of purified proteins with histone peptides that were trimethylated, unmodified, or acetylated at lysine 27. The bar chart shows mean HRP signal (n = 4 replicate ELISAs, bars = SDM).

### Bivalency contributes to Pc-transcription factor binding in an additive manner

To determine whether the contribution of the additional PCD domain is additive or synergistic, we exposed tethered histone peptides to varying concentrations of soluble PcTF and Pc_2_TF. Histone peptides were immobilized on glass slides as spot arrays by conjugation of biotinylated peptides to crosslinked neutravidin. Red fluorescent signal from the recombinant proteins allowed us to detect the amount of bound protein without the aid of a dye-conjugated antibody (Fig. 4A). We detected no interaction with unmodified histone H3 peptides and very little signal over background for the Pc_Δ_TF negative control. We carried out titrations of the recombinant proteins to determine affinities of each protein for 10, 20 and 50 μM of tethered H3K27me3 ligand (Fig. S1). Figure 4B shows representative trials for 20 μM H3K27me3. The calculated affinity of monovalent PcTF for 20 μM of H3K27me3 was 5.14 - 8.95 μM for four independent trials (Fig. 4B and S1). This range of values suggests similar or higher affinity of a fusion-PCD compared to isolated PCD peptides that were analyzed in other *in vitro* experiments. In other work, fluorescence polarization values for ~60 amino acid PCD homologues from Drosophila and mouse were Kd = 5.0 ±1 μM [13] and 165 ±20 μM [12], respectively. Thus, PCD retains its intrinsic binding affinity as an N-terminal motif within a fusion protein. Overall, at 10, 20 μM of ligand the Kd value of Pc_2_TF was roughly 2-fold the value of PcTF (Fig. S1). The highest concentration of ligand showed a 4- to 6-fold increase for the bivalent protein in two trials (Fig. S1).

**Figure 4.**
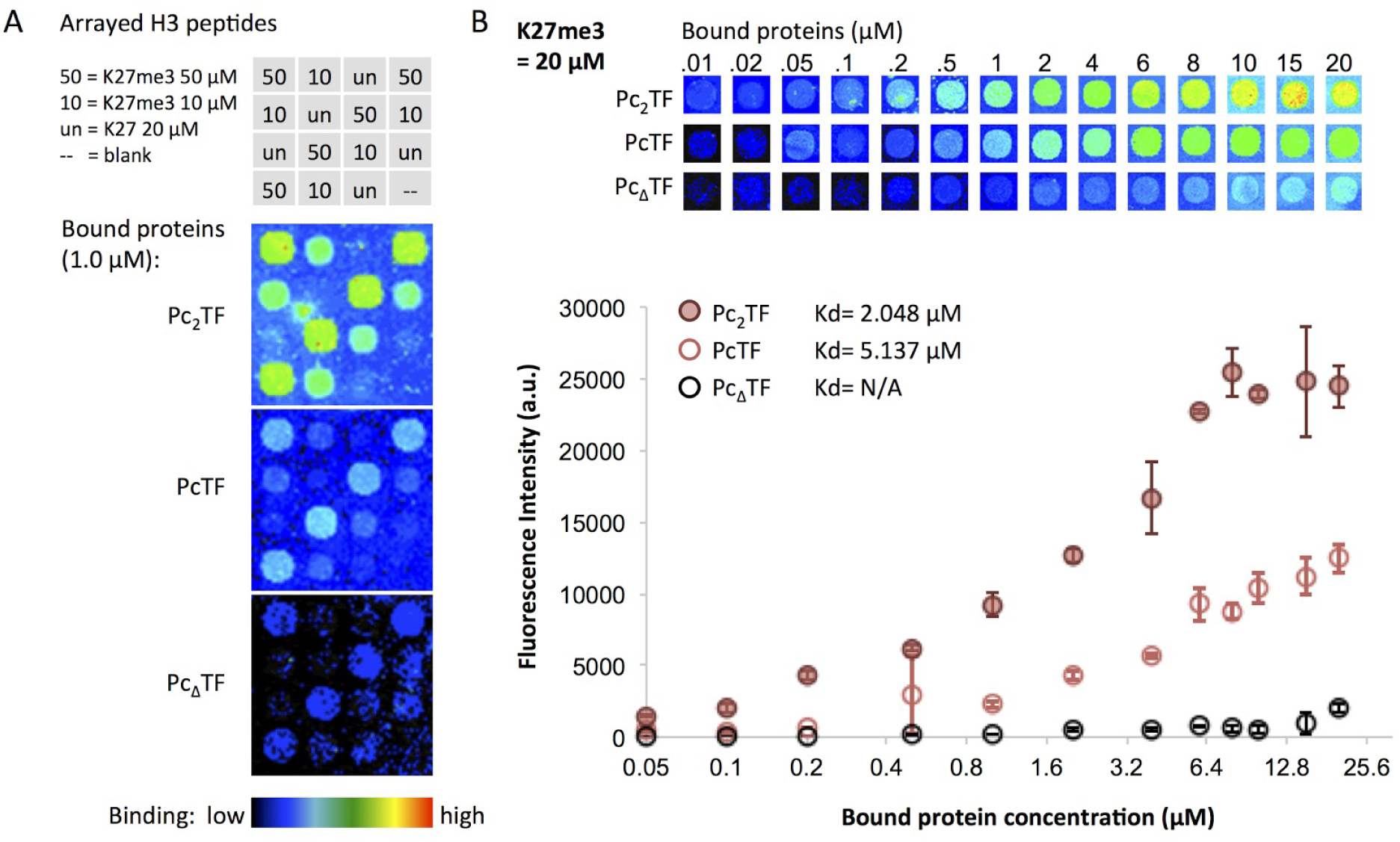
Binding affinities for PcTF and Pc_2_TF were determined using spot array assays. (A) Slides were spotted with histone H3 peptides (K27me3 or unmodified K27) as depicted in the grid (top). (bottom) Pseudo-colored i mages of mCherry fluorescent signal probed with 1 μM individual proteins. (B) Fluorescence signal versus concentration of protein added to the spot array was used to calculate binding affinity (Kd). The fluorescence and Kd values displayed in the graph are from a representative set of trials. For each trial n = 4, bars = SDM.

We can conservatively conclude that doubling of the N-terminal PCD motif increases affinity by 2-fold for the majority the conditions in which we tested soluble proteins (ELISA and spot assay). This result suggests that the additional PCD motif contributes to protein binding in an additive manner *in vitro.* These results raise the question, what is the biological consequence of increased affinity in living cells where the distribution of H3K27me3 is much different? In the cellular chromatin environment, H3K27me3 can occur in *cis* on the radial surface of a single nucleosome (Fig. 1B), in *trans* where DNA bending brings the H3 tails of neighboring nucleosomes close together, or sparsely distributed across many nucleosomes. Furthermore, H3K27me3 marks in living cells are dynamic; the enzyme EZH1/2 adds methyl groups to H3K27, and KDM6A (UTX) and KDM6B (JMJD3) remove these marks (reviewed in [37]). Therefore, we set out to compare Pc_2_TF to PcTF in a cell based system.

### Bivalent Pc_2_TF activates a target gene in partially-silenced chromatin

We determined the biological significance of enhanced affinity by comparing the gene-regulation activities of Pc_2_TF and PcTF at an epigenetically silenced *Tk-luciferase* reporter gene in HEK293 cells. To compare the activities of the fusion proteins in live cells, we cloned the open reading frames for PcTF and Pc_2_TF into a mammalian expression vector called MV10 (Fig. 5A). We observed nuclear RFP signals 48 hours after transfection with Lipofectamine-plasmid DNA complexes (Fig. 5B) and used RFP-positive cell frequency (Table S2) to normalize gene activation in subsequent assays.

**Figure 5.**
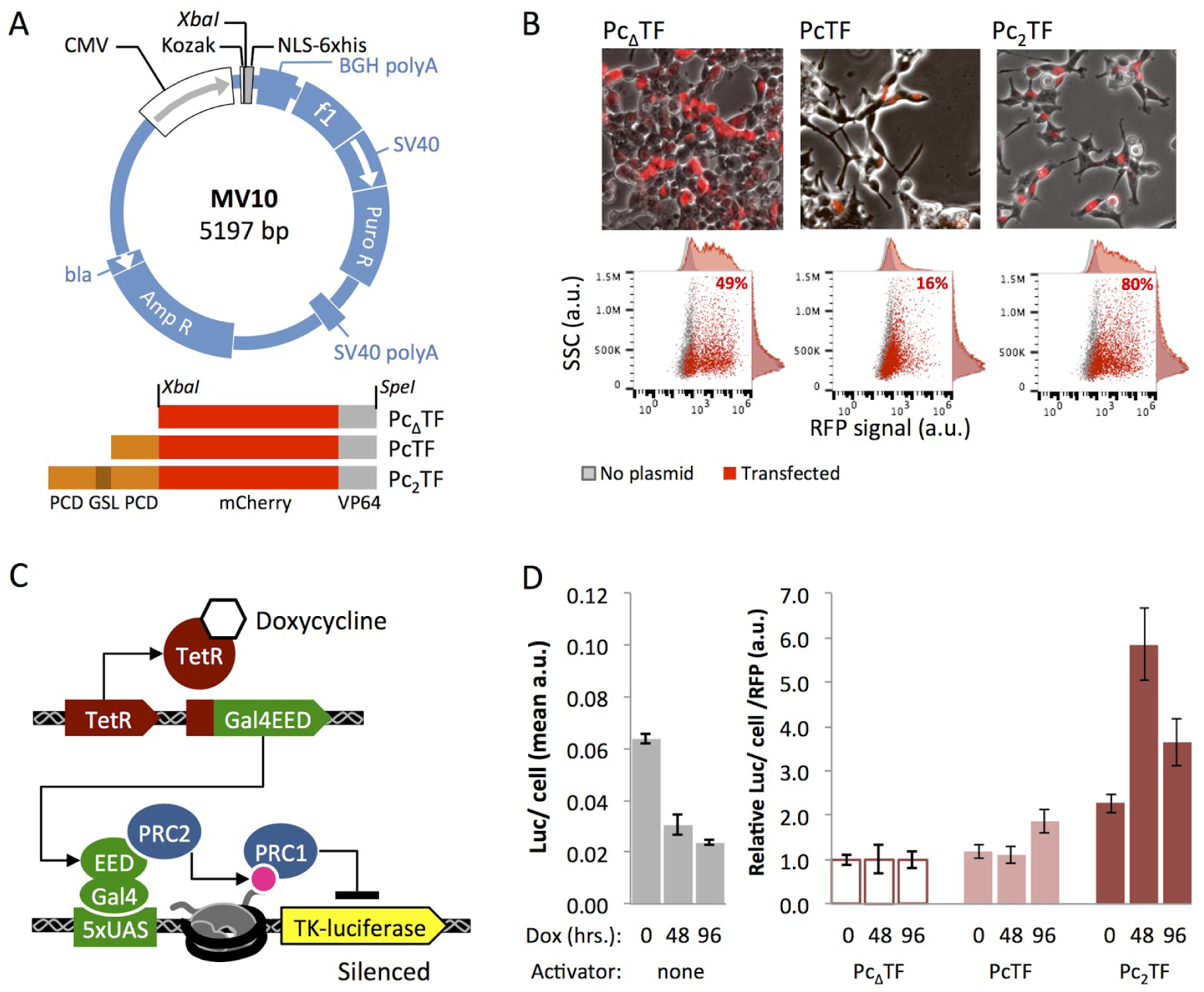
Pc_2_TF stimulates expression at a Polycomb-silenced reporter gene. (A) Fusion constructs were cloned into the MV10 vector at *Xbal.* (B) Fluorescence microscopy confirms nuclear localization of the fusion proteins. Representative examples are shown here. Expression frequency (%RFP-positive live cells) was determined by flow cytometry. Data for dox untreated cells are shown. See Table S2 for all values. (C) An engineered HEK293 cell line, Gal4-EED/luc, was used for doxycycline-mediated control of H3K27me3 and Polycomb-mediated silencing at a *Tk-luciferase* reporter. (D) (left) *Tk-luciferase* expression determined by firefly luciferase assays (n = 3, bars = SDM). Expression is partially silenced prior to dox treatment, as demonstrated previously [39] and becomes fully repressed at 96 hours. Response of Polycomb-silenced *Tk-luciferase* to Pc_Δ_TF, PcTF, and Pc_2_TF is shown in the right bar chart (RFP = RFP-positive cell frequency; n = 3, each scaled to mean Pc_Δ_TF luc/cell/RFP; bars = SDM).

We used a H3K27me3-enriched firefly luciferase reporter gene to compare transcriptional regulation activities of the methyl-histone-binding fusion proteins. HEK293 Gal4-EED/luc cells are engineered to enable doxycycline-mediated control of chromatin state at a chromosomally integrated *firefly luciferase* transgene [38] (Fig. 5C). We and others have demonstrated that doxycycline-induced expression of the Gal4-EED fusion protein leads to accumulation of H3K27me3 near the Gal4 UAS operator and silencing of *Tk-luciferase* expression [10,38,39]. Prior to transfection with fusion protein-expressing plasmids, cells were cultured in 1 ug/mL dox to induce Gal4-EED-mediated silencing at *Tk-luciferase* for 48 or 96 hours, then grown without dox for 48 hours to allow for loss of Gal4-EED from the promoter region. *Tk-luciferase* repression reaches steady state at 96 hours point [39], and repression is maintained by epigenetic inheritance after loss of Gal4-EED [38].

We transfected untreated or dox-treated cells with Pc_Δ_TF, PcTF, or Pc_2_TF. Cells were collected for luciferase activity assays 48 hours after transfection. Pc_Δ_TF did not stimulate activation of *Tk-luciferase* above the negative control background. In cells where silencing was induced for 96 hours, Pc_2_TF showed roughly twofold activation of *Tk-luciferase* compared to PcTF. This twofold relative enhancement of gene regulation activity is consistent with the *in vitro* assays (ELISA and spot arrays). Therefore, the relative binding and affinity observed for purified proteins translates to gene regulation activity in live cells at the model reporter gene tested here.

We also found that Pc_2_TF activates *Tk-luciferase* prior to dox-induced silencing. Previously, we demonstrated that untreated Gal4-EED/luc cells show an intermediate, partially silenced level of *Tk-luciferase* expression [39] compared to fully active *Tk-luciferase* in a “Luc14” parental cell line that lacks the Gal4-EED gene [39]. Furthermore, H3K27me3 was detected via ChIP-qPCR near the *luciferase* promoter (Tk) in uninduced Gal4-EED/luc cells at significantly higher levels than in Luc14 cells. Dox treatment resulted in a further decrease in *Tk-luciferase* expression and a significant increase in H3K27me3 accumulation. In the experiments reported here, basal *Tk-luciferase* expression (Fig. 5D) agrees with independent experiments from our previously reported study (0.02 - 0.07 luciferase activity per cell, a.u.) [39]. The uninduced state may have low levels of H3K27me3 at nucleosomes near the reporter gene in all cells, or high levels of H3K27me3 at the reporter gene in a small proportion of cells in the population. In contrast to Pc_2_TF, monovalent PcTF only activated *Tk-luciferase* after silenced chromatin had been induced for 96 hours. These results suggest that Pc_2_TF is more tolerant of low levels of H3K27me3 in cellular chromatin. In the Conclusions section, we discuss how this idea is consistent with the behavior of the natural bivalent chromatin complex Rpd3S.

## CONCLUSIONS

In our previous work, we have demonstrated the use of a monovalent synthetic effector to activate chromatin-silenced genes in live cells. Natural bivalent chromatin proteins that recognize two histone post translational modifications at once suggest a broader design space for synthetic chromatin effectors. Our application of bivalency to design a synthetic fusion protein produced three important advances for engineering synthetic chromatin effectors. First, we determined that synthetic linkers allow tethered histone PTM-binding peptides to function within the context of a fusion protein *in vitro* and in live cells. Second, we have established that tandem PCDs increase affinity and gene regulation activity by roughly 2-fold compared to a single PCD. Finally, we demonstrated that tolerance for low target PTM levels can be reproduced in a synthetic bivalent effector.

Here, we demonstrated that different synthetic linkers allow tethered histone PTM-binding peptides to bind *in vitro* to varying degrees. We observed weaker binding for the longer flexible linker (80 amino acids) compared to the shorter linker in our ELISA experiment. This result is likely due to lower production of the long flexible linker variant in TXTL. Given that both variants showed binding above background, GGGGS-repeat number may not significantly affect bivalent PCD engagement with H3K27me3 *in vitro.* For the rigid linker-tethered PCDs, only the longer length (80 amino acids) appeared to support binding. Assuming that this variant protein was properly folded, lack of binding over background for the shorter EAAAR-repeat variant could be caused by suboptimal rotation, *i.e. in trans* instead of *in cis,* of the second PCD away from the 2-D binding surface in the ELISA well. Indeed, alpha-helical linkers have been used to control rotational orientation in pairs of tethered histone binding domains [7] and Zinc Finger DNA binding domains [32]. In the context of cellular chromatin where looping and folding occurs, H3K27me3 would not necessarily be constrained to one face of the Pc_2_TF protein. Valuable lessons and perhaps greater Pc_2_TF performance might be acquired by exploring additional linker variants in cells as well as in *vitro.* Such work is beyond the scope of the studies reported here, which accomplished a major step by identifying a functional bivalent synthetic effector protein that specifically interacts with its target H3K27me3.

We have established that tandem PCDs increase affinity *in vitro* and gene regulation activity in live cells by roughly 2-fold compared to a monovalent PCD. This is the first report of intramolecular, synthetic bivalency for a gene-silencing-associated mark (H3K27me3). Prior to our report, studies of multivalent engagement with histones have exclusively focused naturally-occurring, tandem histone-binding motifs [40–42]. The wide distribution of multivalency within bromodomain family [6] and other effector proteins [19] suggests that multivalent engagement has an important, evolutionarily-conserved biological role. Multivalency appears to largely be represented by cell-cycle and gene-activating effectors. Relatively few multivalent proteins that recognize silencing marks have been studied in biophysical detail. Examples include the chromodomains of the Arabidopsis protein CMT3 [43] and the mammalian protein HP1β [44]; as bivalent homodimers, these proteins show enhanced interaction with their respective ligands H3K9meK27me and H3K9me3. Pc_2_TF is novel in its composition of histone-binding motifs: adjacent, identical Polycomb chromodomains within a single peptide. Therefore, its activity *in vitro* and in cells provides new insights into the recognition of histone marks by effector proteins.

In the context of cellular chromatin, Pc_2_TF-mediated activation of a target gene *(Tk-luciferase)* appears to tolerate low levels of H3K27me3, whereas the monovalent variant PcTF required greater accumulation of the mark in order to activate the same target. Our ChIP mapping data [39] confirm that low levels appear at the ectopic H3K27me3 site *(Tk-luciferase)* on average. The limited resolution of ChIP-qPCR on chromatin extracted from a large cell population only allows us to surmise that either a few cells have a high local concentration of H3K27me3 near the *Tk* promoter, or that many cells have sparse H3K27me3 in the same location. Enhanced affinity might allow Pc_2_TF to bind more stably (*i*.*e*., have a reduced off-rate) at low H3K27me3 levels and therefore stimulate *Tk-luciferase* expression at higher frequencies. Additionally, the tandem PCD domains might allow stable docking in regions of sparse H3K27me3. Such is the case for the purported bivalent chromatin modifier complex RpdS3, which contains the H3K36me-binding chromodomain protein Eaf3 and an additional histone-tail-binding motif, the PHD finger of Rco1. RpdS3 binds with very similar affinities to nucleosome pairs (dinucleosomes) in which the H3K36me mark is located on both or just one of the nucleosomes [45]. Direct recognition of H3K36me by the Rco1 PHD has not yet been established, and weak interactions with the histone globular domain are thought to contribute to RpdS3 binding; it is unclear whether RpdS3 is ‘truly’ bivalent. Our observation of Pc_2_TF binding with histone tail peptides in the absence of whole nucleosomes suggests that PTM depletion tolerance in general does not require interactions with histone core regions.

Further engineering efforts to achieve greater, nonlinear enhancement of PcTF/ Pc_2_TF may require changes within the PCD binding motif. The hydrophobic interaction between the methylammonium cage and the methyl-lysine moiety (Fig. 1A) depends upon proper positioning of PCD residues that appear discontinuously in the primary sequence; this positioning requires specific intramolecular contacts of peptide residues within the PCD fold. Reverse engineering and *de novo* design of a new binding pocket through randomization of sequences would likely yield many non-functional proteins. K27-adjacent interactions that support that contribute to interaction with the histone tail [13,23] could be leveraged to enhance affinity. However, increasing the stability by introducing addition hydrogen bonding could overwhelm the hydrophobic, K27me3-specific interaction, and allow PCD to recognize unmodified tails or off-target modifications. Trade-offs between affinity and specificity pose formidable challenges to enhancing PCD affinity. Therefore, the most practical strategy for identifying alternative PCDs is to leverage H3K27me3-specific orthologs and paralogs from various species [46]. It will be important to determine cross-reactivity with various histone modifications since CBX PCD peptides have been shown to bind H3K9me3 [12].

Multivalent engagement of combinatorial histone marks has recently become a key line of evidence to support the controversial histone code hypothesis. Rationally-designed synthetic multivalency will advance this important area of research by exploring functions beyond the limits of pre-existing natural multivalent proteins. Furthermore, engineered chromatin effectors provide a practical tool to support artificial regulation of gene expression states through direct engagement with highly conserved components of chromatin, *i.e.* histone tails and their modifications. Therapeutic, synthetic gene regulators that leverage this mechanism could help circumvent the shortcomings of epigenetic inhibitors, which target chromatin enzymes that are mutated in some cancers [47]. In conclusion, our findings demonstrate that synthetic biology is a powerful tool for fundamental investigations of chromatin biology and epigenetic engineering.

## MATERIALS AND METHODS

### Plasmid Constructs for TXTL and Bacterial Expression

Constructs (Fig. 2A, Fig. 3A) were assembled as BioBrick compatible fragments in vector V0120 [48]. Fragments were PCR-amplified with Phusion polymerase using primers 1-6 (Table S1) and a protocol adapted from New England Biolabs Phusion High Fidelity DNA Polymerase (98°C 0:45, [98°C 0:10, 67°C 0:20, 72°C 0:45] × 25, final extension of 5:00), column purified (Qiagen PCR Cleanup Kit), and double-digested with BamHI and Xhol (Thermo Fisher FastDigest). BamHI/XhoI-digested inserts and 50 - 75 ng BamHI/XhoI linearized pET28(+) vector were ligated at a 3:1 molar ratio in a 20 μL reaction as described in the New England Biolabs (NEB) protocol for T4 ligase (M0202). 5 μL of each ligation was incubated with 50 μL Turbo competent DH5-alpha *E. coli* (NEB) on ice for 5 minutes, transferred to 45°C for 45 seconds, then to ice for 5 minutes, and allowed to recover in 350 μL SOC at 37°C with shaking for 30 minutes. Pelleted cells were resuspended in 50 uL SOC, plated on LB agar (50 ug/mL kanamycin), and grown at 37°C overnight. Colony PCR was performed to identify positive ligation results using primers 6 and 7 (Table S1) and the GoTaq Promega protocol. Plasmids were cloned, extracted (Sigma GeneElute Plasmid Miniprep Kit), and Sanger sequenced for verification prior to protein expression in cell-free TXTL or in *E. coli.* Annotated sequences for all pET28 constructs are available online at Benchling-Hayneslab: Synthetic Chromatin Actuators 2.0 (https://benchling.com/hayneslab/f/rmSYkAAU-synthetic-chromatin-actuators-2-0).

### TXTL: Cell-free Expression

TXTL reactions were set up with the following conditions as previously described [35]: 9 μL lysate, 10 nM final template vector, 0.5 nM σ70-T7 RNA pol vector to a total of 12 μL. A Roche Lightcycler 480 was used to detect mCherry fluorescence with the following: protocol: 30°C for 10 minutes, bring to 29°C for 1 second, scan 533-610 nm, repeat 96 times (total 16 hrs).

### *E. coli* Expression and Purification of Proteins

All selection media contained 50ug/mL kanamycin. Pc_Δ_TF, PcTF, and Pc_2_TF in pET28 were transformed into Rosetta 2pLys DE3 cells and plated on LB agar and grown at 37°C overnight. The next day, a single colony from each was used to inoculate 50 mL LB and grown overnight at 37°C at 300 RPM. The next day, 1 liter of LB in a baffled Erlenmeyer flask was inoculated to an OD600 of 0.1. The cultures were grown to an OD600 = 0.6, induced with IPTG (1 mM final concentration) and allowed to express Pc_Δ_TF and PcTF at 37°C for 5 hours with shaking (220 RPM). Pc_2_TF-expression was carried out overnight at room temperature with shaking (220 RPM) to aid solubility of the protein. Cell disruption and protein purification are described in detail in Supplemental Materials and Methods.

### ELISA Assays

All steps were carried out at room temperature except specifically noted, and all incubations and washes were agitated at 800 RPM on an Eppendorf Thermomixer R. Clear bottom plates (Greiner bio-one #655101) were coated in 50 uL of 20 ng/μL neutravidin in PBS pH. 8.0 overnight at 4 C. The plates were washed the next day 3x with 200 μL 0.2% PBST with 5 minutes of shaking at 800 RPM between washes. The plate was blocked for 30 min at 800 RPM at room temperature with 200 uL 5% BSA in PBST followed by 3x washes of 200 uL 0.2% PBST for 5 minutes each at 800 RPM. 50 μL of 1 uM biotinylated peptides (Anaspec) in 0.2% PBST were incubated at room temperature for 1 hour at 800 RPM, followed by 3x washes of 200 μL 0.2% PBST for 5 minutes at 800 RPM. The plate was then blocked for 30 minutes with 200 μL 5% skim milk in 0.2% PBST at room temperature shaking at 800 RPM. 50 uL of purified proteins (put optimal concentration here) diluted in 5% skim milk in PBST were incubated for 1 hour at room temperature at 800 RPM. The wells were washed 3x with 200 μL 5% skim milk in PBST with 5 minutes of 800 RPM shaking. 100 μL of 1:3000 chicken anti-mcherry (company and product number) in 5% skim milk in 0.2% PBST were incubated for 1 hour at 800 RPM and room temperature followed by 2x of 200 uL 5% skim milk in 0.2% PBST for 5 min each at 800 RPM. 100 uL of 1:3000 Rabbit anti chicken HRP (RCYHRP Genetel 0.5 mg/mL) in 5% skim milk in 0.2% PBST was incubated for 30 minutes at room temperature and 800 RPM. The plate was washed 5x with 200 μL 0.2% PBST for three minutes each at 800 RPM. The plate was incubated with 100 μL of 1-step Ultra TMB-ELISA (Thermo-Fisher #34029) for 15 minutes while blocked from light. The plate was then deactivated with 100 μL 2.0 M sulfuric acid, incubated for 2 minutes, and read at 450 nm.

### Peptide Spot Arrays

APTES functionalized glass slides were coated with 200 μL of 1:1 (v/v) 40 mg/mL BS3 crosslinking solution and 1 mg/mL neutravidin with a cover slide and incubated overnight at 4°C. The next day, the cover slide was removed and the slide was rinsed 3x with 0.2% PBST for 5 minutes each. The slides were deactivated by incubation with Na_2_CO_3_/NaHCO_3_ buffer pH 9.4 for 30 minutes. The slides were quickly rinsed with ddH_2_O, and centrifuged to dry at 1200 RPM for 2 minutes. The slides were printed with biotinylated peptides at concentrations of 10, 20, or 50 μM in 20% glycerol and PBS with a pin-printer and incubated at room temperature for 1 h to allow the biotinylated peptide bind to neutravidin. The distance between spots were 600 um. The slide was rinsed with ddH_2_O as described above and blocked with superblock for 1 hour at room temperature. Proteins were diluted in superblock and incubated on the slide for 1 hour at room temperature. The slides were rinsed with 0.2% PBST for 3 minutes each followed by quick rinsing with ddH2O 3x and centrifuged dry (as described). Red fluorescent protein (mCherry) signal was detected at 50% gain and 50% intensity on a PowerScanner at 635 nm and 535 nm, 10 um resolution. The slides were also scanned at 75%-75% and 100%-100% to to obtain a sutiable signal to noise ratio. Arraypro software was used to quantify the median intensity values for each spot and background levels. Graphpad Prism software was used to fit the data to the binding saturation nonlinear regression equation y=(B_max_*X)/(K_d_+x), where B_max_ is the highest binding value and X is the concentration of protein.

### Plasmid Constructs for Mammalian Expression

MV10 was constructed from pcDNA3.1(+) (Invitrogen) with the following modifications. The CMV promoter was removed via SpeI digestion and T4 ligase recircularization. A dsDNA fragment that encodes *Kozak* (ribosome binding site), *XbaI,* a nuclear localization sequence, 6xhistidine, and a stop codon (5’-cccgccgccaccatggagtctagacccaagaaaaagcgcaaggtacaccatcaccaccatcacgcgtaaagctgag) with *SpeI* overhangs at both ends (ctag/t) was inserted at *XbaI.* CMV (SpeI/XbaI fragment) was reintroduced upstream of Kozak at *SpeI.* Proper orientation of inserts was confirmed by Sanger sequencing. Constructs PcTF and Pc_2_TF (Fig. 5A) were PCR-amplified (Phusion) with primers 9 and 10 (Table S1), double-digested with XbaI and SpeI, and column-purified (Qiagen PCR Purification, 28104). Construct Pc_Δ_TF (Fig. 5A) was double-digested with XbaI and SpeI (Thermo Fisher FastDigest) and isolated by electrophoresis and gel purification. XbaI/SpeI fragments and 25 ng Xbal-linearized, dephosphorylated MV10 vector were ligated at a 2:1 molar ratio in a 10 μL reaction as described in the Roche protocol for the Rapid DNA Ligation Kit (11635379001 Roche), using 1.0 μL NEB T4 ligase instead of the supplied enzyme. All 10 μL of each ligation was incubated with 50 μL Turbo competent DH5-alpha *E. coil*(New England Biolabs) on ice for 5 minutes, transferred to 45°C for 45 seconds, then to ice for 5 minutes. Cells were plated directly on pre-warmed LB agar (100 ug/mL ampicillin) without recovery and grown at 37°C overnight. Plasmid DNA was prepared (Sigma GeneElute Plasmid Miniprep Kit) from 5 mL cultures inoculated with single colonies. Proper orientation of the inserts was determined by XbaI and PstI double-digestion of prepped plasmids and Sanger sequencing. Annotated sequences for all MV10 constructs are available online at Benchling-Hayneslab: Synthetic Chromatin Actuators 2.0 (https://benchling.com/havneslab/f_/rmSYkAAU-svnthetic-chromatin-actuators-2-0).

### Cell Culture and Transfection

HEK293-Gal4-EED cells were grown in Dulbecco's modified Eagle's medium (DMEM) supplemented with 10% tetracycline-free fetal bovine serum and 1% penicillin and streptomycin at 37°C in a humidified CO_2_ incubator. Silencing of the reporter gene *(Tk-luciferase)* was induced by supplementing the media with 1 μg/mL of dox for 48 or 96 hours. For wash-out of doxycycline, growth medium was removed and replaced with dox-minus medium supplemented with 0.5μg/mL puromycin to select for the transgenic anti-Gal4-EED shRNA [38], and grown for 5 days. Prior to transfection, dox treated or untreated cells were plated in 12-well culture dishes at 40% confluency (~1.0E5 cells per well) in 2 mL pen/strep-free growth medium. Transient transfections were carried out by adding 100 μL of DNA/Lipofectamine complexes to each well: 1 μg DNA (10 μL), 3 μL Lipofectamine LTX (Invitrogen), 87 μL Opti-MEM.

### Imaging and Flow Cytometry

48 hours after transfection, cellular mCherry (580/610 excitation/emission) was imaged in culture dishes on a Nikon Eclipse Ti wide field fluorescent microscope (MEA53100, filter G-2E/C). For flow cytometry, the growth medium was removed, adherent cells were gently washed with 1xPBS, harvested by trypsinization (Trypsin-EDTA 0.25%, Gibco), resuspended in growth medium to neutralize the trypsin, pelleted at 200 RCF, and resuspended in 1xPBS supplemented with 1% FBS. Red fluorescent signal from mCherry was detected on a BD Accuri C6 flow cytometer (675 nm LP filter) using CFlow Plus software. Data were further processed using FlowJo 10.0. RFP values for transfected cells are listed in Table S2.

### Luciferase Assays

For the same samples that were used for flow cytometry, cells per 100 μL were determined by flow cytometry (BD Accuri C6). 100 μL of cells or 1xPBS (blank) were incubated with 100 μL of complete luciferase assay reagent as described in the protocol for the Biotium Firefly Luciferase Assay Kit (89138-960) and in previous work [39] in Corning and Costar 96-well Cell Culture Plates, black, clear bottom (Bioexpress). Chemiluminescence was detected using a Synergy H1 Multi-Mode Reader (Biotek). Luciferase expression per cell was calculated as follows: Sample Luciferase per cell = [Sample Luciferase signal] - 1xPBS blank signal/[cell count × (100 μL/20 μL)]. Values for transfected cells were divided by RFP frequency (Table S2).

## ACKNOWLEDGEMENTS

The work was supported by the NIH NCI (K01 CA188164 to K.A.H.). S.T., D.V., and L.S. were supported by the Biological Design PhD Program at ASU. D.V. was also supported by the Western Alliance to Expand Student Opportunities (NSF HRD 1401190). We thank undergraduate researchers B. Laughlin, C. Gardner, D. Tze, and J. Xu for early contributions to plasmid construction. We also thank V. Noireaux for his generous gift of TXTL reagents.

## CONTRIBUTIONS

S.T. built bacterial expression constructs, produced purified protein, performed protein validation, and executed the TXTL experiments. S.T. and L.S. designed and executed ELISA and spot array analyses. D.V. built mammalian expression constructs, and performed transfections, flow cytometry and imaging, dox-mediated silencing, and luciferase activity assays. K.A.H. was responsible for oversight of the experiments and writing of the manuscript.

